# Virus-induced overexpression of heterologous FT for efficient speed breeding in tomato

**DOI:** 10.1101/2023.04.13.536731

**Authors:** Yingtian Deng, Antonia Yarur-Thys, David Baulcombe

## Abstract

Potato Virus X (PVX) vectors expressing the *Arabidopsis thaliana FT* or tomato *FT* ortholog *SINGLE-FLOWER TRUSS* (*SFT*) accelerated tomato flowering time by 14-21d and increased by 2-3 fold the number of flowers and ripe fruit compared with non-infected or empty vector-infected plants. The *Arabidopsis* FT was more effective than the tomato orthologue and there was no alteration of the plant architecture, leaf, flower, or fruit morphology. The virus is not transmitted to the next generation and viral vectors with expression of a heterologous FT will be a useful approach to speed breeding in tomato and other species.

## INTRODUCTION

Tomato is an a widely cultivated South American crop that is also a model species for basic research because it is informative about aspects of growth and development that cannot be addressed using *Arabidopsis* – the standard model in plant biology. It is useful, for example, for investigation of fleshy-fruit formation (Kimura and Sinha 2008) and patterns of branching during vegetative development (Xu et al. 2015). A limitation of tomato, however, is the long generation time. It is typically 65-100d and a system for speed breeding (Wanga et al. 2021) of tomato would, therefore, benefit both basic and applied research.

One approach to speed breeding exploits the FLOWERING LOCUS T (FT) regulator of the floral transition (Kardailsky et al. 1999; Kobayashi et al. 1999; Koornneef 1991). The FT protein is transported through the vasculature to the floral meristem where it acts together with its activator CONSTANS (CO)(An et al. 2004; Corbesier et al. 2007; Takada and Goto 2003). The role of FT as a mobile inducer of flowering is conserved in a wide range of plants (Abelenda et al. 2016; Lin et al. 2007; Tamaki et al. 2007; Krieger et al. 2010) and its transgenic expression stimulates early flowering and reduces generation time (Lewis and Kernodle 2009; Wigge 2011).

An alternative, non-transgenic method of speed breeding uses viruses as expression vectors. For use of such vectors a gene of interest is inserted into the vector at a site that does not interfere with viral genes and it is expressed, often at very high levels, in the infected plant. This virus-mediated overexpression (VOX) allows tracking of virus infection in the plant (Abrahamian et al. 2020) with reporter genes, and it is useful for functional analysis of plant genes (Lee et al. 2012; Lindbo et al. 2001; Manning et al. 2010; Majer et al. 2017; Zhang et al. 2013). VOX of FT results in Virus-Induced flowering (VIF) in crop breeding programs (McGarry et al. 2017) and research. It has advantages over transgenic FT expression because there is no genetic manipulation of the plant genome, and it can be deployed on multiple cultivars without the need for multiple transformations or crossing.

Potato virus X (PVX) is a well-established virus vector with potential in VIF. This vector is infectious on a broad range of *Solanaceous* plants, was initially developed for VOX (Chapman et al. 1992) but has been used widely for virus-induced gene silencing (VIGS) (Lu et al. 2003; Ruiz et al. 1998). PVX-based VOX of FT in tobacco was used in a mutation analysis of FT function (Qin et al. 2017) but, previously it had not been tested as system for speed breeding of tomato.

In this study we used a PVX-vector to test the effect of *Arabidopsis thaliana* and tomato (*Solanum lycopersicum*) FT (AtFT and SlySFT respectively) on flowering and fruiting in tomato. Our findings validate the previously demonstrated use of virus vectors for VOX of FT as an approach to speed breeding. We also illustrate how the overexpression of AtFT was more effective than SlySFT most likely because RA silencing of the endogenous gene is minimised. Our findings confirm the potential for PVX-mediated expression of FT as a convenient and efficient system for speed breeding in tomato and indicate that heterologous FT genes may be more effective than those from the species being infected.

## RESULTS and DISCUSSION

### PVX mediated gene silencing and expression in tomato

To test the PVX system for VIF we used PVX vectors with the intact ORF of FT from tomato and *Arabidopsis*. The tomato and Arabidopsis FT homologues have 73.6% and 69.3% identity at the protein and nucleic acid levels respectively. We also constructed vectors for VIGS with fragments of PDS from tomato and *Nicotiana benthamiana* (*Nb*) cDNA. The *PVX::PDS* constructs would induce VIGS-mediated photobleaching that would report the extent of virus vector movement in the infected plants. All constructs had the inserted sequence placed into the MCS site of the PVX vector *pGR107* (Figure 1a) and they were in Ti plasmid vectors so that *Agrobacterium* infiltration could be used for virus inoculation. Inoculation of *Nb* was a positive control that has been well characterized with the PVX vector system.

**Figure 1.**
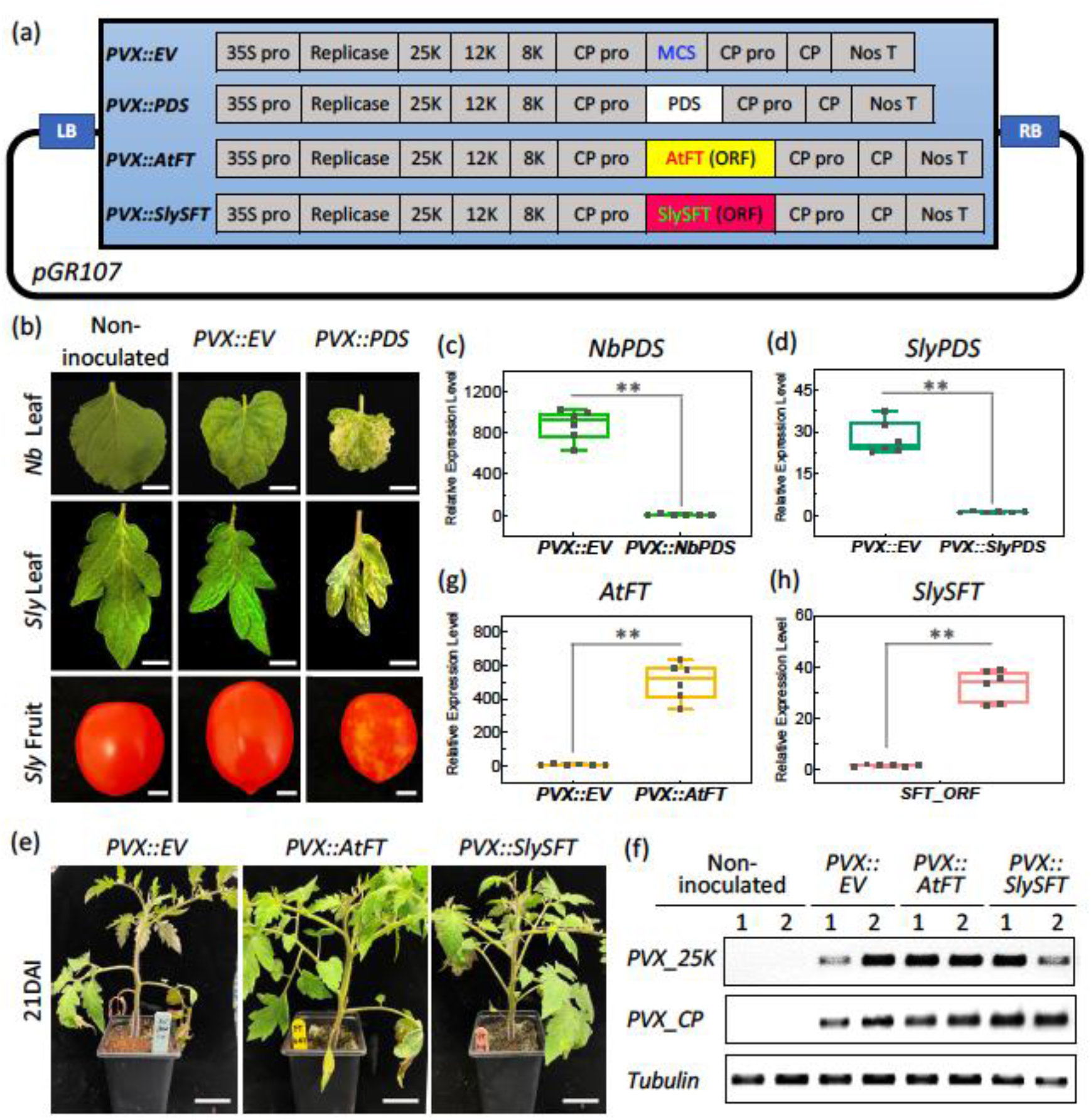
PVX vectors and PVX-induced VIGS and VOX in tomato. (a) Schematic diagram of *PVX* vector components; (b) *PVX*-induced *PDS* silencing phenotypes in *N. benthamiana* leaves, tomato (*Sly*) leaves and fruit, bar = 1cm; (c) and (d) Relative expression level of *PDS* in *PVX:EV-* and *PVX:PDS-*inoculated tobacco (c) and tomato (d) leaves at 21DAI, (**, P < 0.01); (e) *PVX:FT*-inoculated tomato plants at 21DAI, bar = 2cm; (f) Abundance of PVX coat protein and 25K RNA in tomato plants from (e) determined by RT-PCR; (g) Relative expression level of *AtFT* in *PVX:EV* and *PVX:AtFT* treated tomato leaves at 21DAI determined by RT-qPCR, (**, P < 0.01); (h) Relative expression level of *SlySFT* in *PVX:EV* and *PVX:SlySFT* treated tomato leaves at 21DAI determined by RT-qPCR, (**, P < 0.01).

At 21 days after infiltration (DAI) with *PVX::PDS*, but not with the control constructs, there was photobleaching due to silencing of *PDS* in both species (Figure 1b, *Nb* and *Sly* leaves, *PVX::PDS* panels) but not in the controls (Figure 1b, *Nb* and *Sly* leaves, non-inoculated and *PVX::EV* panels). PDS transcript levels assayed by qRT-PCR were lower in the photobleached leaves of the *PVX::PDS* inoculated plants than in the controls (Figure 1c and 1d). There was also photobleaching of tomato fruits by *PVX:PDS* (Figure 1b, *Sly* fruit, *PVX::PDS* panels) indicating that the PVX vector was persistent in the inoculated plants until fruit ripening. These results confirmed that the PVX vector is functional in tomato.

*PVX::AtFT* and *PVX::SlySFT* were inoculated to tomato cotyledons at 14 days after germination (DAG) to investigate the potential of this system for VIF. The inoculated plants developed normally with mild PVX-induced systemic symptoms (Figure 1e) and there was accumulation of PVX RNA (Figure 1f). Furthermore, qRT-PCR analysis confirmed VOX because *AtFT* and *SlySFT* RNA were more abundant in plants inoculated with the respective PVX vectors (Figure 1g, 1h). There was, however, a lower FT over-expression level with the *SlySFT* construct.

### PVX-induced AtFT and SlySFT expression accelerates flowering and fruit production in tomato

Both PVX and PVX-expressed FT would be transported towards the shoot apical meristem (Hein et al. 2005; Jaeger and Wigge 2007) and, consistent with effects on flowering, the inflorescences appeared earlier and the floral buds were more numerous on plants infected with *PVX::AtFT* and *PVX::SlySFT* than on the controls (Figure 2a, pink arrowheads). The open flowers appeared consistently at 42-48 DAG rather than 55DAG (Figure 2b, yellow arrowheads) and the ripe fruit appeared at 80-90DAG rather than 100DAG or longer in the controls (Figure 2b, red arrowheads and Fig.2c). Consistent with the VOX of FT (Fig.1g and h) the acceleration was greater with *PVX::AtFT* rather than *PVX::SlySFT*.

**Figure 2.**
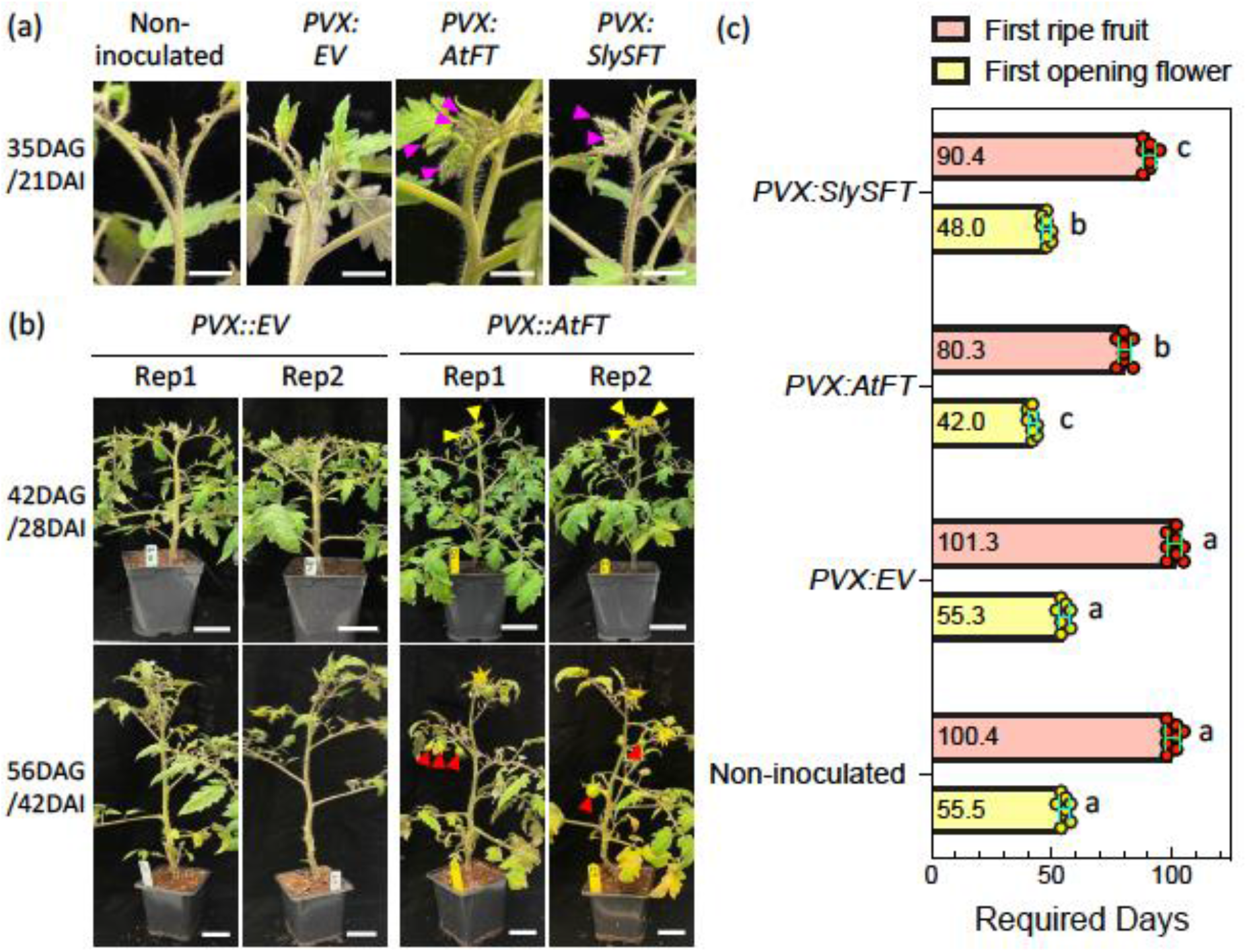
PVX-based VOX of FT accelerated flowering and fruiting in tomato. (a) Apical shoots of non-inoculated and *PVX:EV-, PVX:AtFT-* and *PVX:SlySFT-*treated tomato plants at 21DAI, bars = 1cm; (b) *PVX:EV* and *PVX:AtFT* treated plants at 28DAI and 42DAI, bars = 2cm; (c) Timing of first flower and first ripe fruit on non-infected and *PVX:EV-* infected controls or *PVX:AtFT-PVX:SlySFT-*infected plants, the average value are shown on the bottom of histogram (P < 0.05).

The number of flowers and fruits also increased in the *PVX::AtFT*-and *PVX::SlySFT*-infected tomato plants. At the 42DAI the plants infected with the FT constructs had 100% (*PVX::AtFT*) or 87.5% (*PVX::SlySFT)* more open flowers than the controls (Fig. 3a-h, Fig. 3q). By 126DAI the plants infected with the FT constructs had 100% (*PVX::AtFT*) or 62.5% (*PVX::SlySFT)* more fruit than the controls (Figure 3i-p, Fig. 3q). These findings validate the PVX-induced expression of FT with AtFT proving more effective than SlySFT in all of our assays. We cannot rule out a protein-mediated basis for this difference but, given the difference in FT RNA levels (Fig. 1g and h), a more likely explanation invokes a VIGS effect.

**Figure 3.**
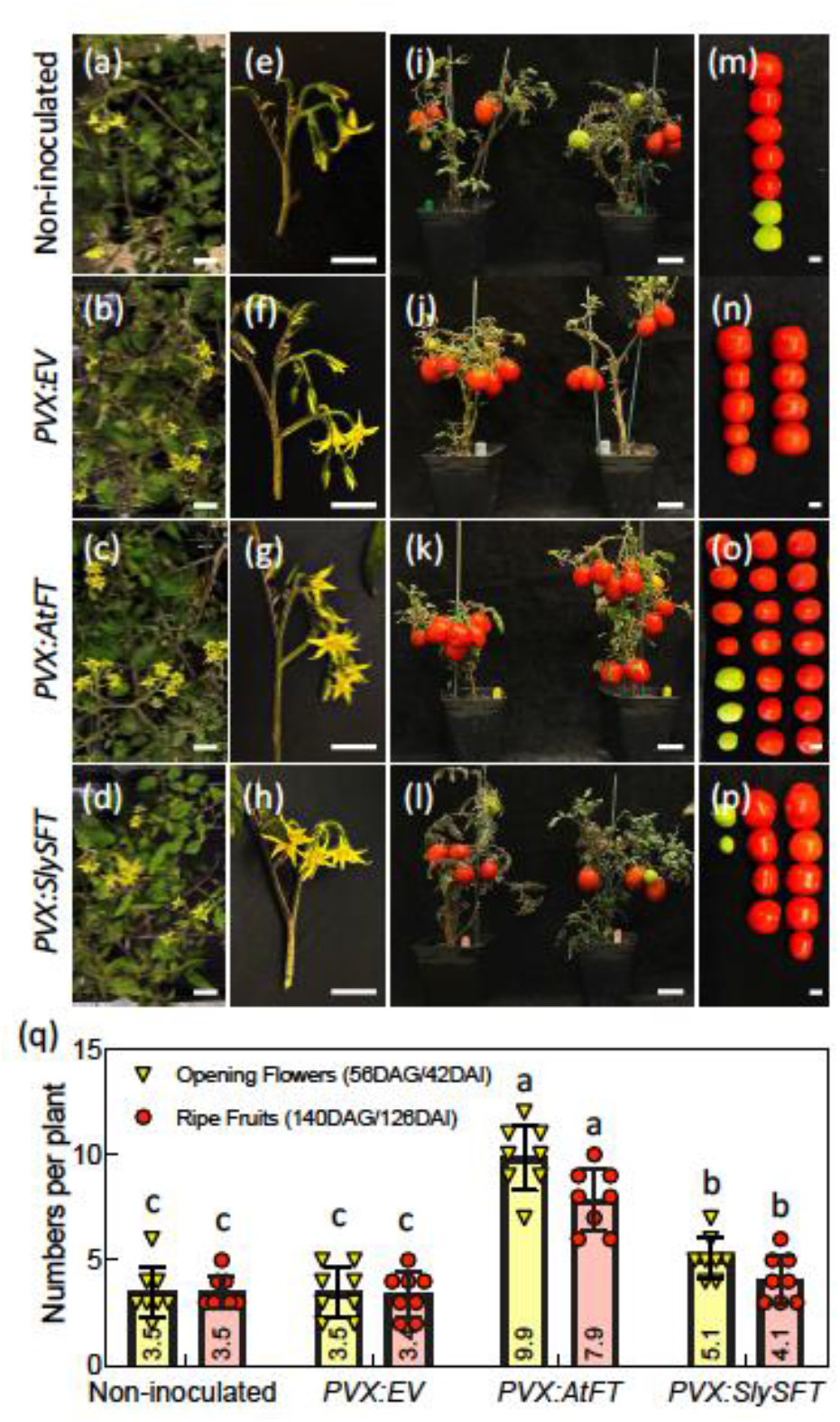
*PVX* induced *FT* expression increase flower numbers and fruit yield in tomato at a time point. (a)-(d) Top view of non-inoculated (a), *PVX:EV* (b), *PVX:AtFT* (c), and *PVX:SlySFT* (d) plants at 42DAI, bars = 2cm; (e)-(h) A single inflorescence of non-inoculated (f), *PVX:EV* (g), *PVX:AtFT* (h), and *PVX:SlySFT* (i) plants at 42DAI, bars = 2cm; (i)-(l) non-inoculated (i), *PVX:EV* (j), *PVX:AtFT* (k), and *PVX:SlySFT* (l) plants at 126DAI, bars = 5cm; (m)-(p) Fruits form plants in (i)-(l), bars = 2cm; (q) Number of opening flowers and ripen fruits in non-inoculated, *PVX:EV, PVX:AtFT* and *PVX:SlySFT* plants at 42DAI and 126DAI stage respectively (P < 0.05).

To explore this possibility, we designed RT-PCR primers from the ORF and 3’ UTR that would amplify cDNA corresponding to the endogenous rather than viral FT RNA (Figure 4a). The results showed that *SlySFT* endogenous transcript decreased in *PVX::SlySFT*-inoculated leaves more than with *PVX:EV*-or *PVX:AtFT* (Figure 4b). It is therefore likely that VIGS of the endogenous *FT* RNA in *PVX::SlySFT*-inoculated leaves resulted in lower levels of FT than with *PVX:AtFT*.

**Figure 4.**
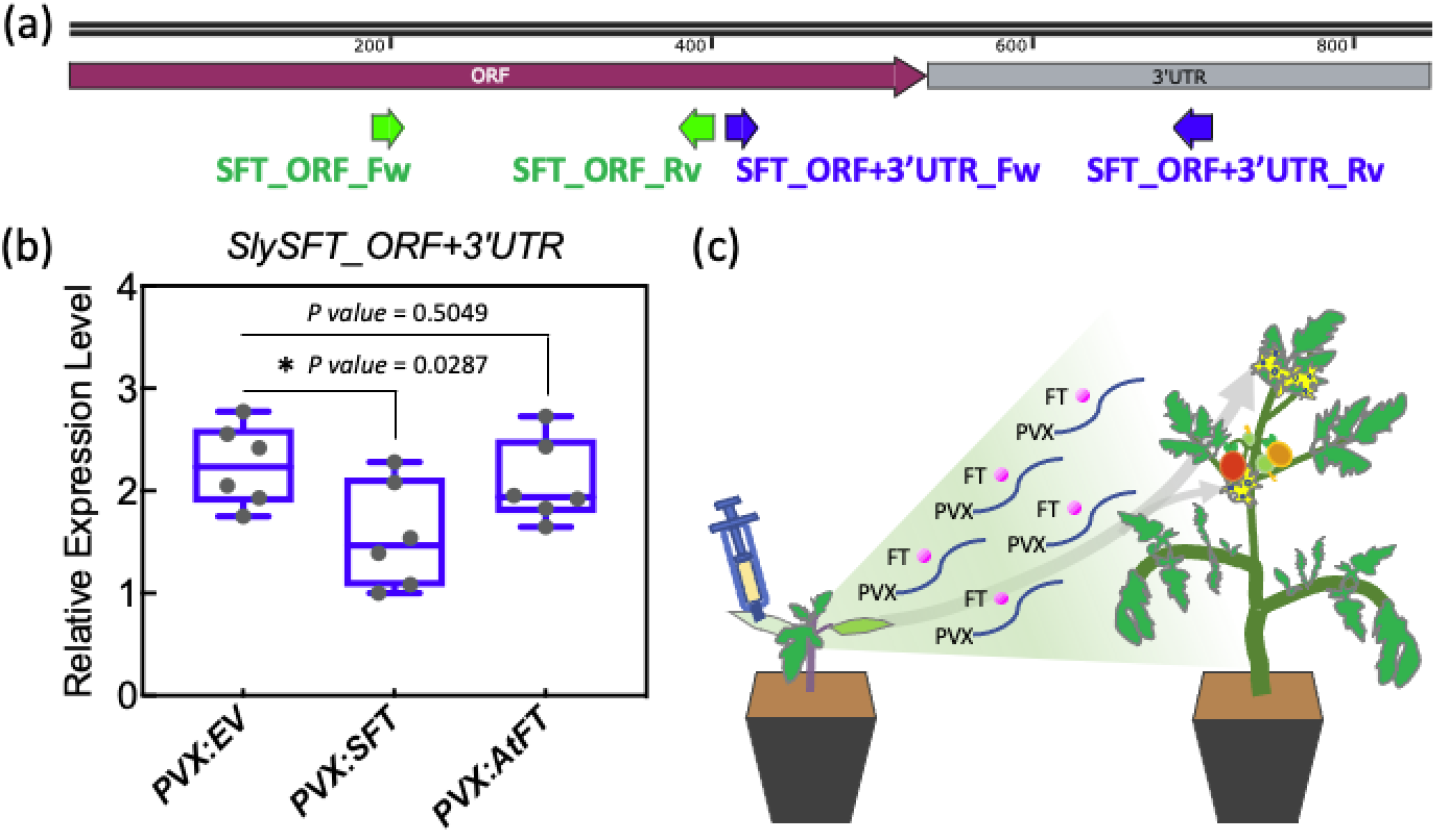
Endogenous *SFT* gene in the host tomato plant is partially silenced when expressing *PVX:SlySFT*. (a) Schematic diagram of *SlySFT* cDNA sequence, and the primers used for testing expression level of over-expressing SFT from PVX vector (green) and endogenous *SFT* gene in the host tomato plant (blue) (*, P < 0.05); (b) Relative expression level of *SlySFT_3’UTR* in *PVX:EV-, PVX:SlySFT-* and *PVX:AtFT-*inoculated tomato leaves at 21DAI determined by RT-qPCR; (c) Working model of PVX induced FT expression.

The VOX effect did not persist into the next generation. Seedling progeny of the infected plants (Figure 5a) did not have viral symptoms or detectable viral RNA and we are confident that VIF-mediated speed breeding could be used in tomato without the possibility of vertical transmission of the PVX vector.

**Figure 5.**
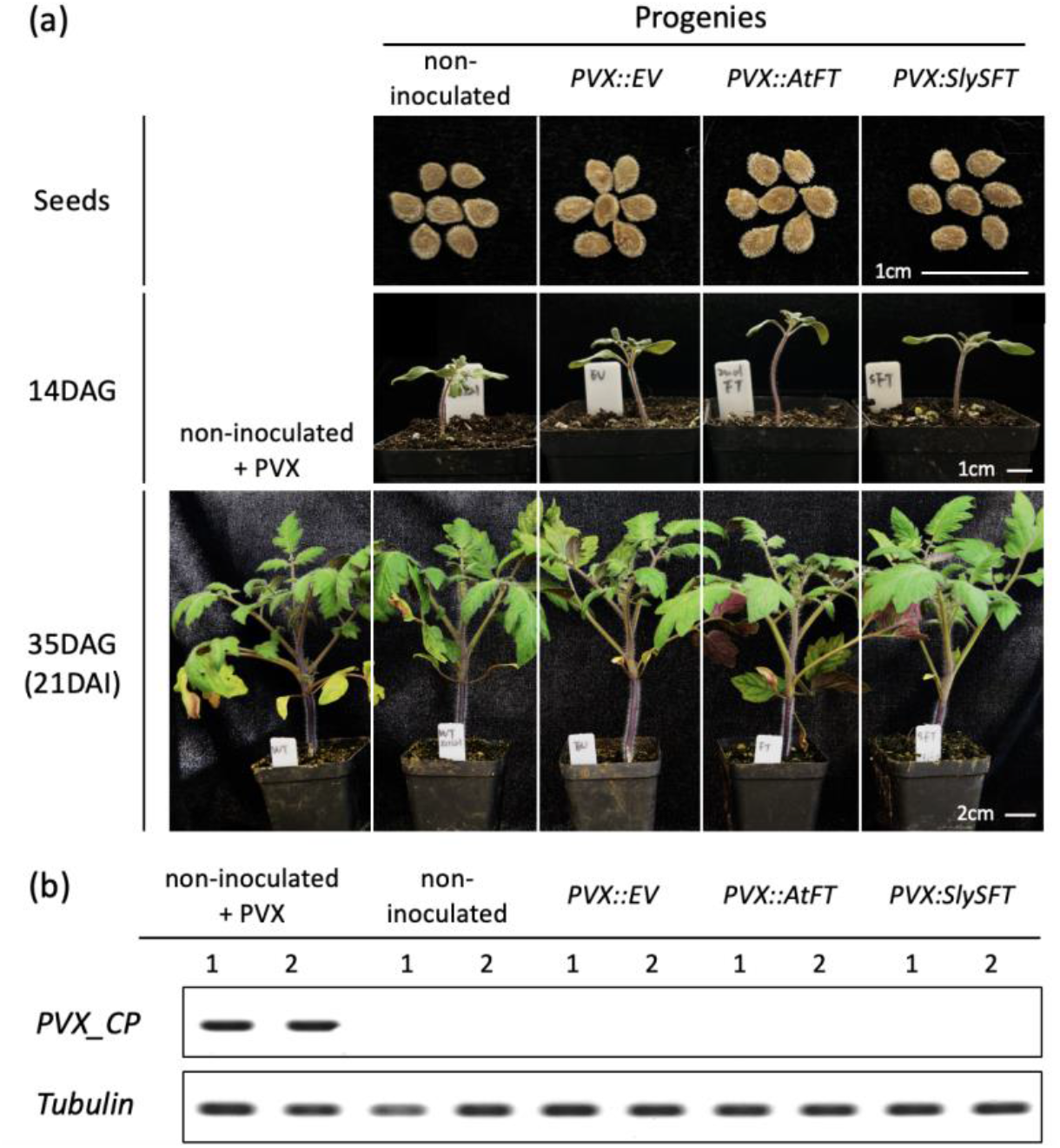
No PVX virus was detected in the progeny of PVX-infected tomato plants. (a) Seeds (top), 14DAG seedlings (middle), and 35DAG plants (bottom) of inoculated and non-inoculated tomato plants, with a group of non-inoculated progenies being inoculated with PVX for 21 days (bottom and left); (b) Abundance of PVX coat protein of tomato plants in (a, bottom) determined by RT-PCR.

Our findings advance previous technology in two ways. First they validate the use of PVX as an alternative to the TRV vector system for use in tomato (Brigneti et al 2004). PVX has a mono-rather than bi-partite viral RNA genome and may be simpler to use in many contexts. A second advance is the reduced endogenous RNA silencing resulting from the use of a heterologous gene for over-expression (Figure 4). This strategy to avoid RNA silencing of endogenous genes had a big effect on the overexpression of *FT* (Figures 2 and 3) may also be generally relevant when using virus vectors for gene overexpression.

## Materials and Methods

### Plant materials and growth conditions

Tomato ‘M82’ (Solanum lycopersicum) plants were grown in the plant growth facility under long-day conditions (light/dark:16h/8h) at a temperature of 25°C before and after inoculation.

### PVX vectors recombination

The PVX constitutive expression vector pGR107 was firstly generated and kept in our lab (Lu et al. 2003). For PVX vectors recombination, 325bp fragments of the *SlyPDS* (Solyc03g123760.3.1, ITAG4.0), full length of *SlySFT* (534bp, Solyc03g063100.2.1, ITAG4.0) and *AtFT* (AT1G65480, 528bp, Araport11) was PCR-amplified from tomato or Arabidopsis cDNA, and the PCR product was inserted into *SmaI*-cut pGR107vector. Sequences and primers used in these constructions are listed in the Supplementary Files.

### Tomato inoculation

*Agrobacterium* strains contain the above empty or recombined vectors were cultured overnight in LB liquid medium. The bacterial cells were collected by centrifugation and resuspended in infiltration medium (IM, 10 mM MES, 200 μM acetosyringone). *Agrobacterium* cells were then resuspended in an optical density at an OD600 of 0.1. The *Agrobacterium* suspensions were infiltrated into two-weeks-old tomato cotyledon (14DAG) by using a needleless syringe.

### RNA extraction and Reverse transcription PCR (RT-PCR) analysis

Total RNA was extracted from tomato leaves at 21DAI with Trizol (Invitrogen™, USA). First-strand cDNA was synthesized using 2 μg total RNA with random primer by using RevertAid First Strand cDNA Synthesis Kit (Invitrogen™, USA). RT-PCR was performed using the DreamTaq Green DNA Polymerase (Thermo Scientific™, USA) and the PCR products were running in the 1.2% agarose gel. Primers used in the RT-PCR are listed in the Supplementary Files.

### Reverse transcription - Quantitative PCR (qRT-PCR) analysis

By using the above reverse transcribed cDNAs as templates, qRT-PCR was performed using the Luna^®^ Universal qPCR Master Mix (NEB, USA) in a 10 μL total sample volume (5 μL 2× Luna^®^ Universal qPCR Master Mix, 1.0 μL of primers, 1.0 μL of cDNA, and 3 μL of distilled, deionized water). Relative gene expression values were calculated using the 2^−^ΔΔ^Ct^ method. The tomato *Tubulin* gene (Solyc04g081490.3.1, ITAG4.0) and *SKP1* gene (Solyc01g111650.3.1, ITAG4.0) were used as internal reference genes. At least three biological replicates were included for each treat, and each replicate was from independent sampling. Primers used in the RT-qPCR are listed in the Supplementary Files.

### Data analysis and software

All the statistical analyses in this study were performed by using Student t-test method. The histograms and boxes graphs were generated via GraphPad Prism 9.5.0 (GraphPad Software, San Diego, CA, USA).

## Acknowledgements

We thank Ms Mel Steer (Department of Plant science, Cambridge university) for helping tomato horticulture.

